# Pubertal development and hypothalamic-pituitary-gonadal axis are altered in male mice lacking *Mecp2*

**DOI:** 10.1101/2025.08.19.671012

**Authors:** Ana Martín-Sánchez, Daniela Jiménez-Díaz, Rafael Esteve-Pérez, Alexandru Vasile-Tudorache, Jordan E. Read, Sasha R. Howard, Carmen Agustín-Pavón

**Author notes:** Equal contribution (Daniela Jiménez-Díaz, Rafael Esteve-Pérez). Corresponding authors: **Ana Martín-Sánchez**,**, Carmen Agustín-Pavón,**.

## Abstract

**Background:** Mutations in the *MECP2* gene, encoding the epigenetic reader Methyl-CpG binding protein 2, are the main cause of Rett syndrome, a rare neurodevelopmental disorder. Besides severe symptoms such as profound intellectual disability, loss of speech and motor skills and epilepsy, loss of function of *MECP2* has been associated with pubertal dysregulation, but the biological mechanisms leading to this remain unclear.

**Methods:** Using a mouse model of Rett, in which males are hemizygous and females heterozygous for *Mecp2* loss of function mutation, we assessed the onset and progression of puberty, together with increase in body weight and onset of neurological symptoms in post-weaning mice until puberty. In brain samples of young adult mice, we analysed hypothalamic Gonadotropin releasing hormone (GnRH) neurons by immunofluorescent labelling, and in plasma samples measured circulating GnRH, LH and testosterone concentrations. Finally, we analysed testosterone dependent arginine-vasopressin circuits.

**Results:** In our mouse model we found delayed puberty in *Mecp2 ^CD1^*-null males, associated with a reduced rate of weight gain, but with puberty onset occurring at a lower body weight than in wildtype controls. Despite later puberty onset, *Mecp2^CD1^*-null male mice were found to have an increased number of GnRH neurons, but displayed lower levels of circulating reproductive hormones. Consequently, *Mecp2^CD1^-*null males have deficient testosterone-dependent arginine-vasopressin innervation. In female *Mecp2^CD1^*-heterozygous mice, we found no overall significant differences in pubertal development or GnRH neurons. The lack of significant alterations in females might be related to a later onset of neurological symptoms due to heterozygosity.

**Conclusions:** Our data supports that *MECP2* is essential for typical pubertal development, with complete loss of *Mecp2* in a male murine model resulting in abnormalities of pubertal timing related to lower body weight, with an observed increase in hypothalamic GnRH neurons.

## Introduction

The X-linked *MECP2* gene encodes the methyl CpG-binding protein 2 (MeCP2), a multifunctional protein first characterised as a transcriptional repressor able to bind to methylated DNA (1) and currently mostly defined as an epigenetic reader with both transcriptionally repressive and activating functions (2) MeCP2 is expressed in a variety of tissues, but it has a fundamental role in the brain, where it is involved in the maintenance of a mature neuronal phenotype (3,4). Thus, mutations in *MECP2* are associated with several neurological disorders (5). In particular, loss of function mutations in *MECP2* are the main cause of Rett syndrome (RTT) (6), a rare neurodevelopmental disorder and one of the most severe conditions linked to this gene. RTT mainly affects girls, who are heterozygous for the mutation; on the contrary, boys suffer a complete loss of *MECP2* and typically die perinatally (5). Girls with RTT, who are born apparently healthy, develop the first symptoms around 6-18 months of age, including an arrest of development followed by loss of acquired skills. The main symptoms are loss of speech, loss of purposeful use of the hands, intellectual disability, epilepsy and progressive loss of ambulation (7). On the other hand, duplication of *MECP2,* causing duplication syndrome (MDS), affects boys, leading also to intellectual disability, motor impairment and epilepsy (7).

In addition to these severe symptoms, loss of function of *MECP2* can lead to endocrine dysregulation, which can contribute to emotional symptoms (8), and disordered puberty (9). Related to this, a handful of studies have reported cases of patients with RTT or MDS that display precocious puberty (10–12). Recently, rare variants of *MECP2* were identified in patients with central precocious puberty with isolated neurodevelopmental features, including autism or microcephaly, but without RTT (13,14). Further, pubertal trajectories have been seen to be altered in a subset of patients with RTT, with around 20% of females experiencing precocious onset of puberty but delayed menarche (9). This extended course of puberty, with early breast development but late pubertal completion, is also observed in disrupted oestrogen signalling such as from exposure to estrogenic endocrine-disrupting chemicals (15).

The onset of puberty is triggered by the activation of hypothalamic neurons containing Gonadotropin Realising Hormone (GnRH) (16). These neurons are neuroendocrine cells that develop within the nasal placode and reside in the olfactory bulbs, septum and the preoptic and anterior regions of the hypothalamus in the adult mammalian brain (17). GnRH hypothalamic neurons, via pulsatile GnRH signalling, regulate the gonadotrope cells of the hypophysis, which release luteinizing hormone (LH) and follicle-stimulating hormone (FSH). In turn, LH and FSH control the production and release of gonadal hormones (oestradiol, testosterone), which provide negative (and positive) feedback signalling to the hypothalamus-pituitary to coordinate pubertal development and adult reproduction.

Several potential pathways exist via which *MECP2* could influence GnRH activation. Firstly, by means of its transcriptional control of FXYD domain-containing transport regulator 1 (*FXYD1*), a protein responsible for control of GnRH neuronal excitability. Patients with RTT and *Mecp2*-null mice demonstrate a pathogenic overexpression of *FXYD1* in the frontal cortex of the brain, attributed to lack of repression by *MECP2* (18), and mRNA levels of *Fxyd1* in rat brain closely correlate with increased GnRH neuron excitability at puberty and first oestrus (19). Next, as an epigenetic regulator, MeCP2 regulates gene expression through interaction with both 5-methylcytosine (5mC) and 5-hydroxymethylcytosine (5hmC) residues in DNA, impacting chromatin accessibility, recruiting histone modifying enzymes and forming part of various polycomb complexes (20–22). Beyond the traditional idea of MeCP2 as a repressor of gene expression through its association with 5mC, it has also been found to associate with active chromatin regions, enriched at 5hmC *loci* in neurons. The Rett syndrome *MECP2* variant p.R133C preferentially disrupts the association with 5hmC (22). Likewise, MeCP2 has demonstrated a strong co-localisation within the mouse cortex with repressing histone mark H3K27Me3 (21) as well as an association with the activating histone mark H3K4Me3 (20), indicating a role as both an activator and repressor of gene expression. Additionally, evidence for estrogen receptor α and β (Erα and Erβ) expression in murine GnRH neurons suggests a direct role for estrogen signalling in the regulation of GnRH secretion (23). MeCP2 is seen to associate with the Erα promoter in mouse cortex, correlating with increased promoter methylation and reduced Erα expression beyond postnatal day 10 (P10). Female mutant *Mecp2* mice demonstrate an increase in this Erα expression beyond P10, inverse to the reduced levels seen in wildtype mice, suggesting a mis-regulation of Erα expression upon loss of functional *Mecp2* (24).

*Mecp2*-mutant mice are valuable model systems for the study of RTT (25). Loss of function mutation is not immediately lethal to males, so *Mecp2*-null male mice survive into young adulthood, albeit developing severe motor impairment and reduced lifespan (25), and early onset of symptoms (26) The onset of symptoms in *Mecp2*-heterozygous females is variable, with some of them displaying mild behavioural symptoms at young age and others remaining pre-symptomatic until 4-6 months of age (8,27). Aside from the typical motor symptoms, in the first report describing the model, Guy and collaborators (25) noted that *Mecp2*-null males had universally undescended testicles and were infertile. However, it is unclear whether males are unable to mate because of their motor impairment, whether they exhibit inappropriate sexual behaviour or display hormonal deficits. Young adult *Mecp2*-null males displayed low levels of testosterone-dependent features, such as aggression, arginine-vasopressin (AVP) sexually dimorphic central circuits and production of male pheromones (28). These data strongly suggest that *Mecp2-*mutant mice could help to understand the mechanisms of altered pubertal development observed in patients, but, to our knowledge, this has not been previously investigated.

Thus, in this study, we sought to characterize the pubertal trajectories, analyse GnRH neuronal circuitry and the levels of circulating sex hormones in a relatively novel model of *Mecp2* mutant mice on CD1 background (29).

## Material and methods

### Animals and rearing conditions

For this study, we crossed *Mecp2*-heterozygous females bred in-house (B6.129P2(C)-Mecp2^tm1.1Bird/J^, The Jackson Laboratory) with pure CD1 males (Crl:CD1(ICR), Charles River Laboratories) for more than 10 generations, to ensure complete derivation to the CD1 strain, as per (29). Our decision to derive the model to CD1 background was in order to maximize colony productivity while maintaining the disease phenotype (27). Moreover, a relatively recent systematic review found that, contrary to conventional assumptions, the use of outbred mice, such as CD1, might be preferrable to the use of inbred strains for biomedical research (30) Animals were housed with *ad libitum* water and food in a room maintained at 22 ± 1°C, humidity 55 ± 10% and a12:12 h light/dark cycle, with lights on at 08:00h. All experimental procedures were approved by the Committee of Ethics and Animal Experimentation of the Universitat de València and treated according to the European Union Council Directive of June 3rd, 2010 (6106/1/10 REV1) and under an animal-usage license issued by the Direcció General de Producció Agrària i Ramaderia de la Generalitat Valenciana (2022/VSC PEA/0288).

### Genotyping and expression of MeCP2 in the brain

For genotyping, we obtained ear biopsies at weaning and after DNA extraction we applied the protocol supplied for this strain by the Jackson Laboratory. To further assess the loss of MeCP2 protein expression in the new strain, we carried out an immunofluorescent detection of MeCP2 in a WT and *Mecp2^CD1^*-null males (Supplementary material, Figure S1).

### Assessment of reproductive capacity of Mecp2^CD1^-het females

To assess whether *Mecp2^CD1^*-het females displayed typical breeding, we recorded the time lapse in days since pairing with a stud male until first delivery, and the number of pups surviving until weaning age (postnatal day 23) in the first litter, at F0, F1, F2, F3 in the original *Mecp2^Bird^*-het strain (n=17), and at the F0, F1, F2, F3, and F9, F10 and >F10 in the *Mecp2^CD1^*-het females (n=28). Breeding pairs consisted in one-two young adult *Mecp2*-het females (7-8 weeks of age) plus one stud male from the pure C56/BL6J (Bird strain) or CD1 (CD1 strain) background.

### Weight gain, puberty onset and oestrous cycle monitoring

All mice were weaned at P23. Mice were weighed daily the first week after weaning, from P25 to P31, to monitor weight gain due to its influence on the onset of puberty (females, WT, n= 16, *Mecp2^CD1^*-het, n=11; males: WT, n=16; *Mecp2^CD1^*-null, n=21). Mice that achieved vaginal opening or balanopreputial separation after P31 were weighed daily until the day that puberty parameter was achieved. Puberty was determined by the day of vaginal opening in females and balanopreputial separation in males, respectively, from postnatal day 25 (P25) (females, WT, n = 23, *Mecp2^CD1^*-het, n = 16; males: WT, n =16; *Mecp2^CD1^*-null, n =21) (31,32). Balanopreputial separation in males was assessed by gently attempting manual retraction of the prepuce. Additionally, we determined the occurrence of the first oestrous in a subset of those females by collecting vaginal smears from the day of vaginal opening until first oestrous (WT, n = 10; *Mecp2^CD1^*-het, n = 10), and oestrous cyclicity in a subset of the females by monitoring the duration of the phases of the oestrus cycle for 10 days after P35 (WT, n = 12; *Mecp2^CD1^*-het, n = 8). Briefly, vaginal cells were collected by gently flushing the external vaginal area with a small amount (50-100 μl) of saline solution (NaCl 0.9%, Braun), using a pipette inserted into a sterile latex bulb. The liquid was slowly released into the vaginal opening and then drawn back into the pipette using the bulb. The process was repeated 4-5 times using the same solution until the resulting fluid became turbid. Then, the solution with cell suspension was dropped over a glass slide, dried using a hot plate, and counterstained using a toluidine blue (Sigma-Aldrich) 0.25% solution (33).

### Perfusion, fixation and tissue sectioning

At two months old, an age considered in mice as young adulthood with full reproductive capacity (34), animals (females, WT, n = 6, *Mecp2^CD1^*-het, n = 6; males: WT, n = 5-6; *Mecp2^CD1^*-null, n = 6-7) were deeply anaesthetized using a dolethal overdose (i.p. injection of 0.02 mg/g of body weight of pentobarbital-based solution). Then, animals were euthanized by transcardial perfusion of phosphate saline solution 0.1 M (PBS 0.1M, pH 7.4) using a peristaltic pump (5.5 ml/min for 2 min) followed by 4% paraformaldehyde in 0.1 M PBS, pH 7.4 (same flux for 5 min). Brains were carefully removed and immediately post-fixed in the same fixative solution overnight at 4°C. Then, brains were cryoprotected (30% sucrose solution in 0.1 M PBS, pH 7.6, 4°C) and cut into five sets of 40µm-thick coronal sections using a freezing microtome (Leica SM 2010R) and stored with 30% sucrose and 0.02% sodium azide in 0.1 M PB at –80°C. In another subset of mice (females, WT, n = 3, *Mecp2^CD1^*-het, n = 3; males: WT, n=5; *Mecp2^CD1^*-null, n=5), gonads were removed, postfixed in PFA 4% and stored with 70% ethanol solution, included in paraffin, sectioned in 10 µm-thick sections and counterstained with haematoxylin-eosin.

### Immunofluorescence for GnRH and AVP

We performed GnRH immunofluorescence in one of the five parallel coronal series from both sexes and genotypes (WT females n= 6; Mecp2CD1-het n = 6; WT males n = 5, Mecp2CD1-null n = 7), and AVP immunofluorescence in another brain series, only from males, from both genotypes (WT n = 6, *Mecp2^CD1^*-null n = 6). Briefly, free-floating sections were washed with 0.05 M TRIS buffered saline pH 7.6 (TBS) (3 × 10 min). Then, sections were: (i) pre-incubated in 3% normal donkey serum (NDS; Sigma-Aldrich, G9023) in TBS with 0.2% Triton X-100, at RT for 1 h, to block non-specific labelling; (ii) incubated with TBS with 0.3% Triton X-100, 2% NDS for 48 h at 4 °C with monoclonal rabbit anti-GnRH primary antibody (1:5000, Invitrogen, AB1567) or rabbit anti-vasopressin IgG (1:1000; Chemicon, AB1565); (iii) incubated with fluorescent-labelled RedTM-X-conjugated at a 1:500 dilution (Jackson ImmunoResearch, 711-295-152) secondary antibody (90 min at RT) diluted in TBS. To reveal the cytoarchitecture in brain sections, they were counterstained prior to mounting by 5 min washes in 4’, 6– diamino-2-feniindol (DAPI, a nuclear staining) at a 1:10000 dilution. After each step, sections were washed 3 times for 5 min in TBS except between step (ii) and (iii). Finally, sections were rinsed thoroughly in TB, mounted onto gelatinised slides and cover-slipped with fluorescence mounting medium FluorSave Reagent (Sigma-Aldrich, 345789).

### Determination of circulating hormonal levels

Blood samples from young adult male mice (WT, n = 5-15; *Mecp2^CD1^*-null, n = 9-13) were obtained from the aortic arch immediately before perfusion (between 10:00 a.m. and 13:00 a.m.) and under anaesthesia conditions. These samples were collected into heparinized tubes, which were rapidly centrifuged at 20.000 g for 15 min at RT (Eppendorf 5424 Centrifuge), until plasma was separated from blood cells. Supernatant (100 μl) was collected and immediately stored at –80C until used. Samples selected for ELISA were processed according to the protocol supplied by the ELISA kit manufacturer for testosterone (Bio-Techne R&D Systems, S.L.U. KGE010), LH (Fine Test, EM1188) and GnRH (ElabScience, E-EL-0071) determination. Optical density was read at 450 nm in a Thermo Scientific Multiskan FC automatic spectrophotometer. Concentrations were calculated using their standard curves.

### Microscopy and image acquisition

Brain and gonadal samples were analysed using a microscope equipped with light and fluorescent lamps (Leica Microsystems series 140269 LEITZ DMRB microscope, digital camera LEICA DFC495). Fiji-ImageJ software was employed to manually quantify the number of follicles and corpora lutea, spermatids and Leydig cells, GnRH-positive and AVP-ergic cells using the *cell counter* plugging. In gonadal samples, number of elements are normalized by the area of the sample, and in brain samples this number is relative to the number of slices containing the area o interest. For AVP-ergic innervation, we quantified the area fraction occupied by AVP-immunoreactive pixels in the image after appropriate thresholding. No manipulations of individual image elements were carried out.

### Statistical analysis

Data were analysed using IBM SPSS Statistics 23.0 and GraphPad Prism. We first checked the data for normality (Kolmogorov-Smirnov’s test) and homoscedasticity (Levene’s test). We performed Student’s t-test, one-way ANOVAs and ANOVA for repeated measurements to analyse the weight variation across different ages. Kaplan-Meier analysis was used to evaluate differences in the observation of vaginal opening, first oestrous in females and balanopreputial separation in males. When applicable, pairwise comparisons were analysed with Bonferroni’s correction. When data were not normally distributed, Mann-Whitney’s test was used for the number of follicles data. Statistical differences were considered significant when p < 0.05.

## Results

### *Mecp2^CD1^*-het females provide larger litters than *Mecp2^Bird^*-het females, while maintaining the neurological phenotype

We first analysed whether reproductive capacity was increased in the *Mecp2*^CD1^-het females, as previously reported (35). In the number of days from pairing with a stud male until delivery of the first litter we found no significant differences between strains (Mann-Whitey test, p > 0.05, Supplementary material, Figure S2A). By contrast, the number of pups surviving until weaning in the first litter was significantly higher in the *Mecp2*^CD1^-het as compared to *Mecp2*^Bird^-het females (Mann-Whitney test, p < 0.01, Supplementary material, Figure S2B).

### Pubertal development of *Mecp2^CD1^*-het females is not significantly different from WT controls

The weight of female *Mecp2^CD1^*-het mice increased across days during the first week post-weaning (repeated measures ANOVA, F_3,23_ = 3.95 p < 0.001, Figure 1a), with no differences compared to WT controls. Similarly, no significant differences in timing of vaginal opening (Kaplan-Meier analysis, p > 0.05, Figure 1b) or weight at first oestrous (Student’s t-test, p > 0.05, Figure 1c) were observed between *Mecp2^CD1^*-het females and their WT counterparts. Further, we found no significant differences between genotypes in the occurrence of first oestrous, that happened 2.8±0.7 days after vaginal opening in WT and 2.6±0.7 days after vaginal opening in *Mecp2^CD1^*-het females (Kaplan-Meier analysis, p > 0.05, Figure 1d), cycle length and oestrous cyclicity for ten days from P35 to P45 (Student’s t-test, p > 0.05 in both cases, Figure 2e, f, and Supplementary Figure S3). Finally, histological analysis of the ovaries revealed the presence of follicles in all phases of development, with no significant differences in the number of follicles in any phase (Mann-Whitney rank test, p > 0.05, Figure 1g) and no histological abnormalities (Figure 1h and h’).

**Figure 1.**
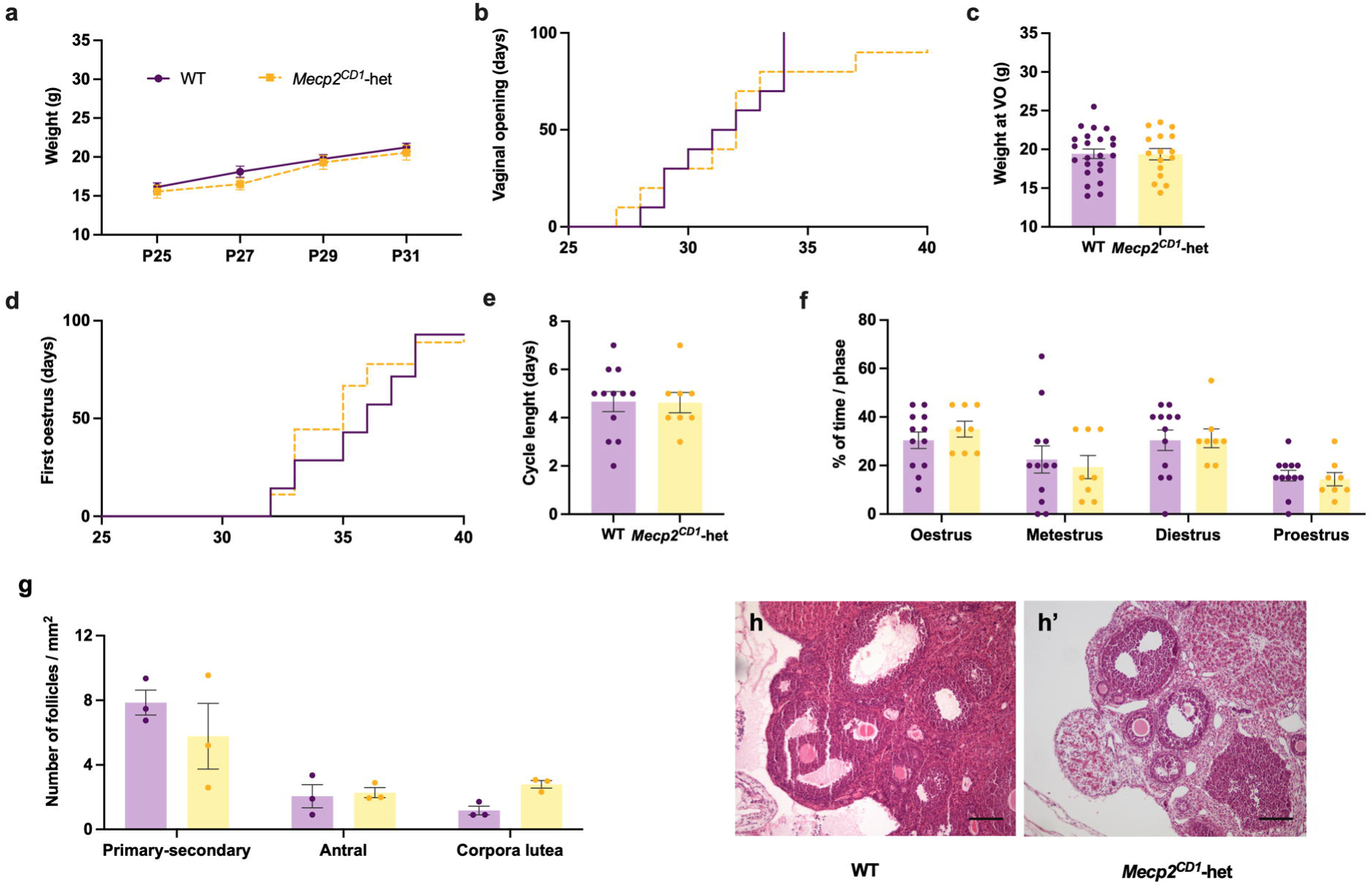
Weight gain, pubertal development, and histology of the ovaries are not significantly affected in *Mecp2^CD1^*-het female mice. a) Graphical representation of body weight gain in *Mecp2^CD1^*-het females in comparison with their WT control mice. b) Days to vaginal opening. c) Weight at vaginal opening. d) Days to first oestrous. e) Cycle length (oestrous to oestrous) and (f) percentage of time in each phase of the oestrous cycle across 10 days of monitoring. g) Number of follicles in different stages. f) Histological images of representative WT (h) and *Mecp2^CD1^*-het (h’) ovaries. Data are shown as Mean ± S.E.M. Scale bar: 50μm

**Figure 2.**
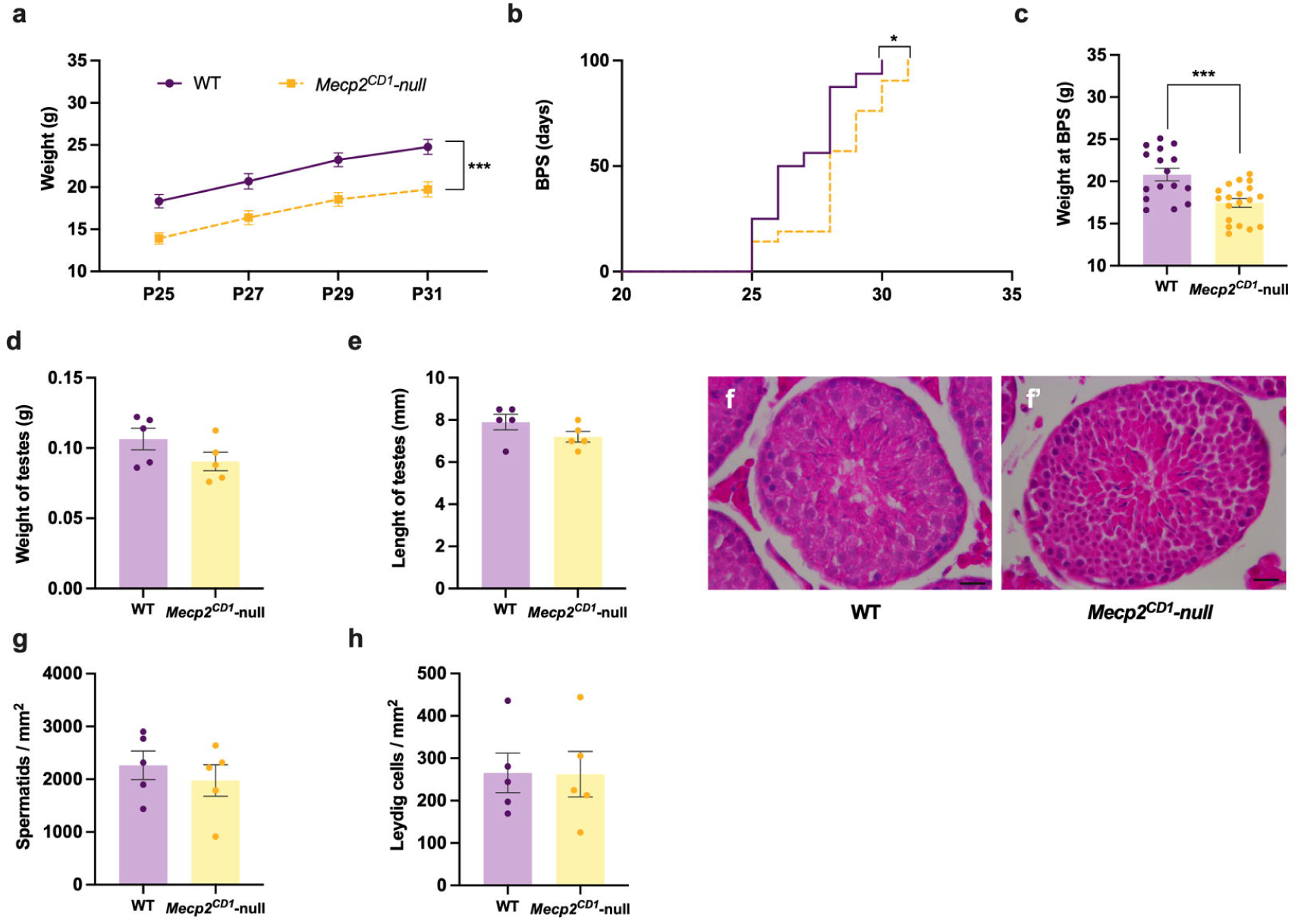
Weight gain and pubertal development in *Mecp2^CD1^*-null male mice are significantly affected. a) Graphical representation of body weight gain in *Mecp2^CD1^*-null males in comparison with their WT control mice. b) Days to balanopreputial separation (BPS) in WT and *Mecp2^CD1^*-null males and c) weight at BPS. d) Length and e) weight of testes. Histological representative images of WT (f) and *Mecp2^CD1^*-null (f’) testes. g) Number of spermatides. h) Number of Leydig cells. Data are shown as Mean ± S.E.M. * p < 0.05; *** p < 0.001. Scale bar: 20μm

### Puberty is significantly delayed in *Mecp2^CD1^*-null males and associated with lower weight

The weight of *Mecp2^CD1^*-null males was significantly lower during the first week post-weaning as compared to WT males (repeated measures ANOVA, F_1,35_ = 15.76, p < 0.001, Figure 2a). The onset of puberty of *Mecp2*-null males showed a significant delay when compared to WT animals (Kaplan-Meier, χ^2^ = 6.025, df = 1, p < 0.05, Figure 2b). Further, balanopreputial separation occurs in *Mecp2^CD1^*-null males at a lower body weight than WT controls (t_33_ = 3.83, p < 0.001; Figure 2b). Of note, and unlike the *Mecp2*-null males on B6.129 background, the testes of young adult *Mecp2^CD1^-*null males were descended and visible, and showed no significant differences in weight (Student’s t-test, p > 0.05, Figure 2d) or length (Student’s t-test, p > 0.05, Figure 2e). Similar to in females, we did not find histological abnormalities in the gonads of pubescent male mice (Figure 2f), observing similar amount of spermatides (Figure 2g) and Leydig cells (Figure 2h) in both genotypes (Student’s t-test, p > 0.05 in both cases).

### The density of GnRH neurons is increased in young adult *Mecp2^CD1^*-null males

Subfertility and aberrant sexual development had been previously linked to the early loss of GnRH neurons in a mouse model of Trisomy 21 (36). Hence, we analysed the distribution and density of GnRH neurons in young adult mice. We found a low number of GnRH-immunoreactive (ir) neurons in the olfactory bulbs (Figure 3a), with no quantitative differences between genotypes in any sex (Figure 3b, c). As expected, the majority of GnRH neurons were mainly located in the septal area and the rostral hypothalamus (Bregma 0.5 to 0.02 mm) (37), both in males and females and in both genotypes (Figure 3d). Further, we corroborated that GnRH neurons co-expressed MeCP2 in WT mice (Supplementary Figure S4a), but, as expected, some GnRH neurons did not express this protein in *Mecp2^CD1^-*het females (Supplementary Figure S4b). Quantitatively, and in agreement with the lack of effect on puberty onset of *Mecp2* mutation in heterozygosity in females, the density of GnRH neurons was not significantly different between *Mecp2^CD1^-*het females and their WT controls (Student’s t-test, p > 0.05, p > 0.05, Figure 3b and e) either in olfactory bulbs or in septo-hypothalamic levels. In contrast, we found an increase in GnRH neurons in *Mecp2^CD1^*-null as compared to WT male mice (t_10_ = 2.83; p = 0.018, Figure 3f) specifically in the septo-hypothalamic area.

**Figure 3.**
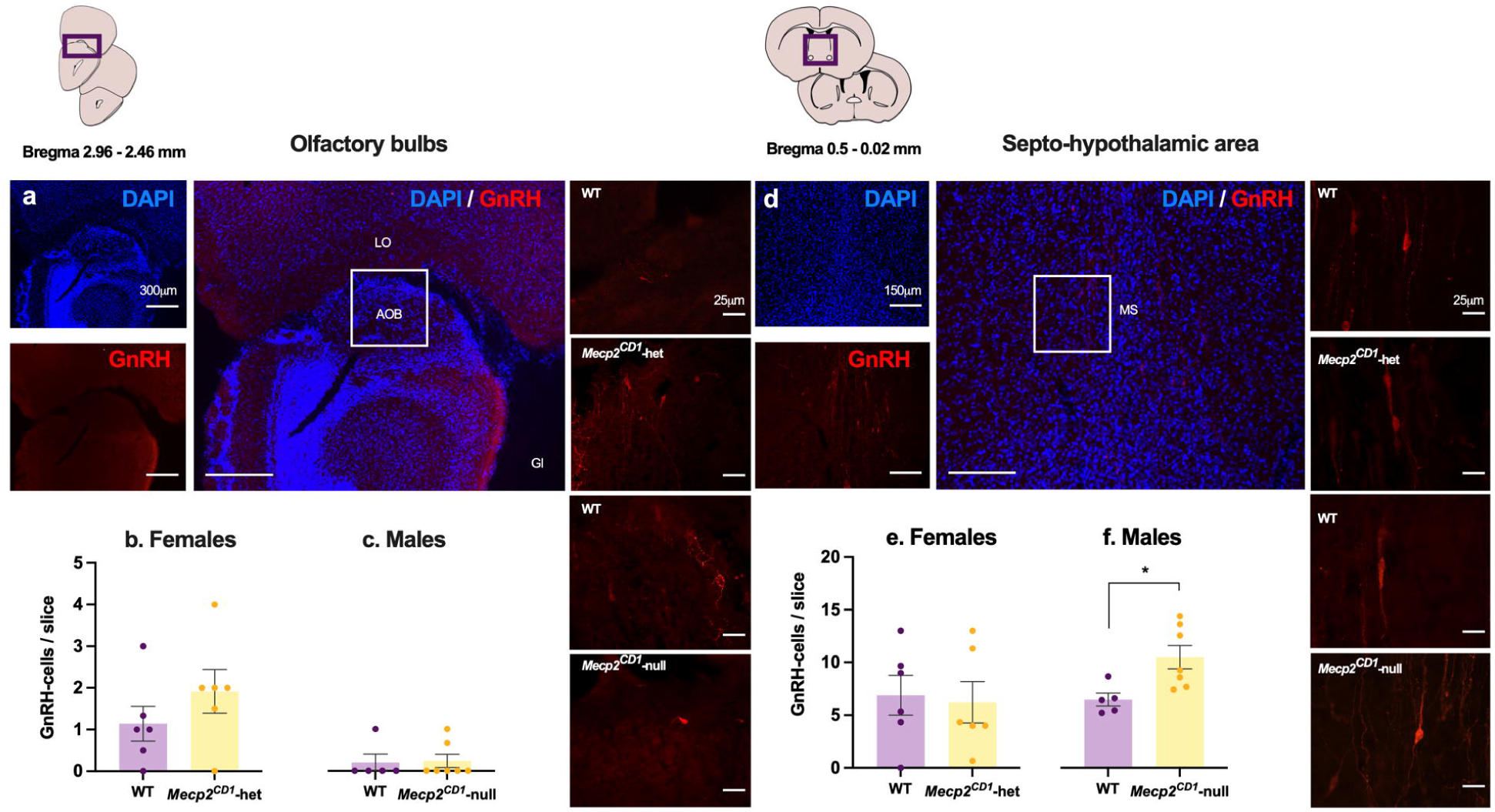
Septo-hypothalamic GnRH-positive cells are increased in *Mecp2^CD1^*-null male mice. a) Representative images of GnRH immunofluoresecence in the olfactory bulbs, where these neurons were scarce both in females (b) and males (c), with no differences between genotypes. d) Representative images of GnRH immunofluorescence in the septo-hypothalamic area. Density of GnRH neurons was not statistically different between WT and *Mecp2^CD1^*-het females (e), whereas the number of GnRH neurons was significantly increased in mutant males as compared to WT controls (f). Data are shown as Mean ± S.E.M. * p < 0.05. Scale bar: 50μm

### Circulating levels of reproductive hormones are reduced in young adult *Mecp2^CD1^*-null males

Given the lack of significant effects of *Mecp2* in heterozygosity in females, we decided to focus further experiments in males. We sought to analyse whether the excess of GnRH neurons in *Mecp2^CD1^*-null males could lead to increased levels of circulating hormones. Surprisingly, we found that relative circulating levels of GnRH showed a trend toward reduction in *Mecp2^CD1^*-null as compared to WT males (t_12_ = 2.08; p = 0.06; Figure 4a). Further, relative LH and testosterone levels were significantly lower in *Mecp2^CD1^*-null males compared to controls (Student’s t test, t_26_ = 2.17, Figure 4b; t_21_ = 2.33, Figure 4c, p < 0.05 in both cases).

**Figure 4.**
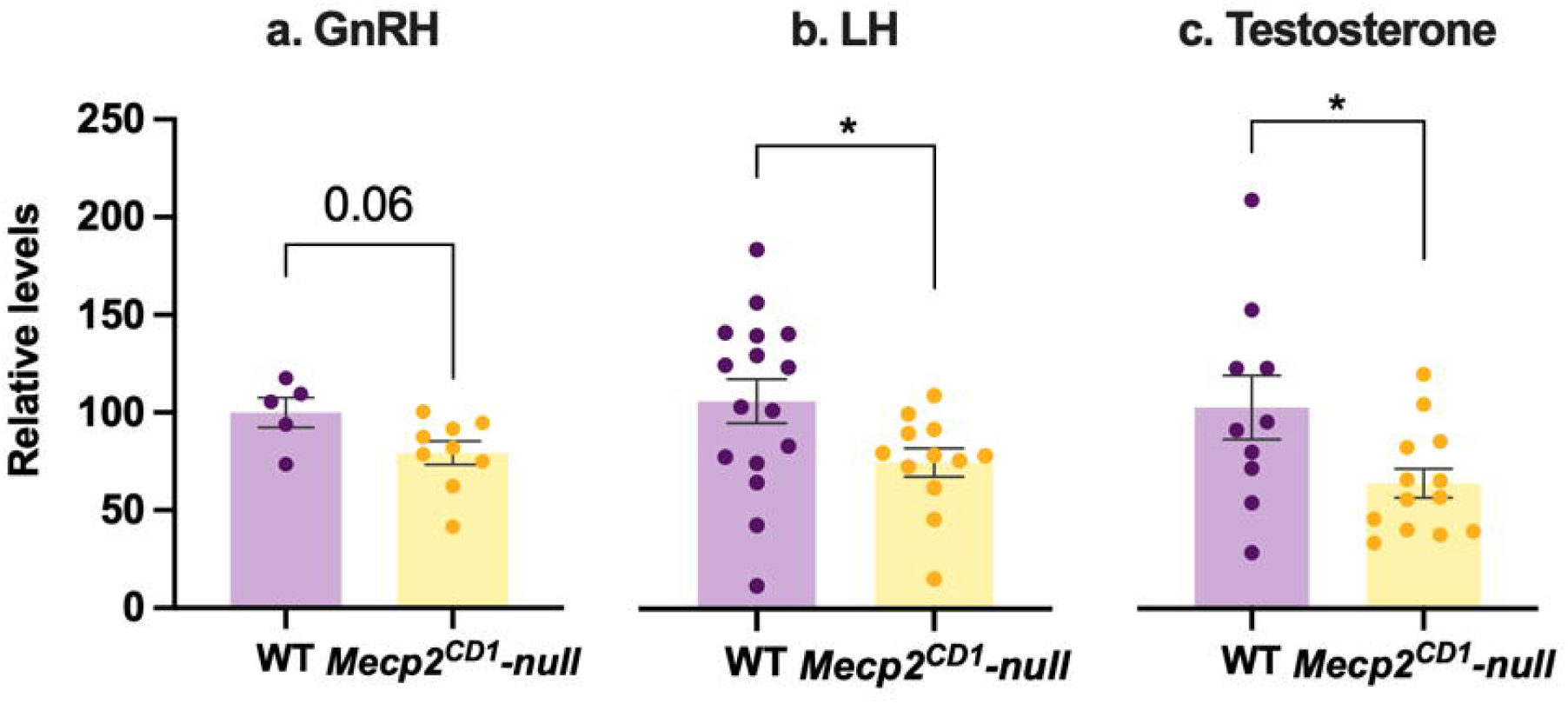
Plasma levels of reproductive hormones in male mice. a) GnRH; b) LH and c) testosterone circulating levels were decreased in *Mecp2^CD1^*-null male mice relative to WT. Data are shown as Mean ± S.E.M. * p < 0.05

### Testosterone-dependent AVP-ergic innervation in the lateral habenula is reduced in young adult *Mecp2^CD1^*-null males

In a previous study using *Mecp2*-null males on B6.129 background, we found significantly decreased testosterone-dependent AVP-ergic innervation in those mice (28). Here, we replicated this measure in *Mecp2^CD1^*-null males, where AVP-ergic innervation was almost absent in the lateral habenula of mutant males (Student’s t-test, t_3_ = 3.02; p < 0.05, Figure 5d, e and f and Supplementary Figure S5a, b and b’). As in our previous study, this reduction appears to be specific to testosterone-dependent innervation, since the number of AVP cells in the paraventricular nucleus of the hypothalamus is similar in *Mecp2^CD1^*-null males and controls, thus not affected by genotype (Student’s t-test, p > 0.05; Figure 5 a, d, c; and Supplementary Figure S5c, d and d’).

**Figure 5.**
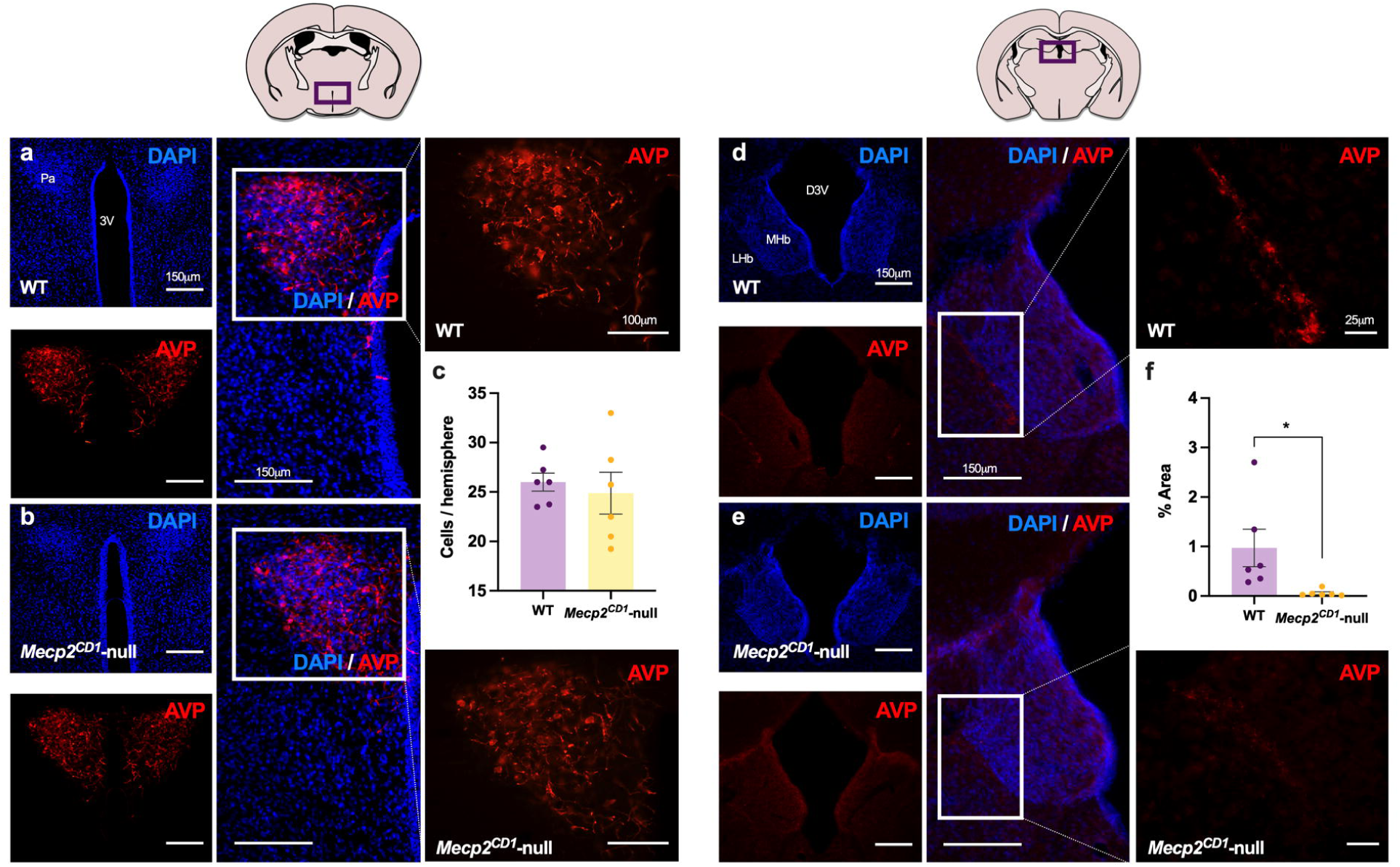
Vasopressinergic testosterone-dependent innervation is reduced in adult *Mecp2^CD1^*-null males. Representative images of the paraventricular nucleus of the hypothalamus (Pa, Bregma –0.58mm) in a WT (a) and a *Mecp2^CD1^*-null male (b). The density of AVP-ergic neurons is not affected by genotype (c). By contrast, representative images of the testosterone-dependent innervation of the lateral habenula (LHb, Bregma –1.58mm) in a WT (d) and *Mecp2^CD1^*-null male (e) show a strongly reduced density of neuronal fibres in mutant animals (f). Data are shown as Mean ± S.E.M. * p < 0.05. Scale bar: 50μm

## Discussion

In this work, we sought to study pubertal dysregulation in RTT syndrome using a mouse model deficient for *Mecp2*, derived to the CD1 strain to improve reproductive capacity. Whereas in heterozygous females we did not find significant effects on pubertal development, we found a significant delay in achieving pubertal milestones in *Mecp2^CD1^*-null males as compared to their WT counterparts, a delay that may be influenced by the significantly lower weight during infancy and pubescence in these males. Further, we found an excess of GnRH neurons in the septo-hypothalamic region in young adult *Mecp2^CD1^*-null males, contrasting with lower circulating levels of LH and testosterone, suggesting a dysregulation of the hypothalamic-pituitary-gonadal axis.

Although RTT syndrome is defined as a neurodevelopmental disorder, it is in fact a multiorgan disease affecting multiple systems in the body, causing gastrointestinal and orthopaedic problems, altered immune response and endocrine comorbidities (38–40). The most common symptoms related to endocrine dysfunction are low bone mineral content, malnutrition (which could be also related to motor impairment causing difficulties in eating) and alterations in the initiation of puberty (9,38,40,41). With regards to puberty timing, data are scarce and conflicting. Various longitudinal population-based studies suggest that a subset of patients with RTT (3-6%) experience premature menarche (9,42), particularly those with milder mutations in the *MECP2* gene (9). In contrast, 19% of girls were found to experience a delayed menarche (9). Overall, data from human patients with RTT suggest that the disease is associated with deviations from normal pubertal timing, with some patients experiencing early puberty onset, premature thelarche but delayed menarche (9).

Body weight and nutrition have a strong influence on pubertal timing and reproductive health (43), and both patients with RTT and the mouse model studied here are likely to be affected by these factors. Indeed, *Mecp2^CD1^*-null males showed a significantly lower weight than healthy controls at all stages, and a significantly delayed balanopreputial separation, which occurred at lower body weight, in comparison to WT males.

By contrast, in our cohort of *Mecp2^CD1^*-het females, body weight was, overall, not significantly different from WT controls, and neither were vaginal opening or first oestrous, a measure which more accurately reflects the onset of human puberty.

Thus, early onset of the disease in males, in which the complete loss of *Mecp2* strongly affects most phenotypes (28,44) and reduces the lifespan of these mice (25), producing low weight and ill health, might have a profound influence on pubertal development in mutant males. Since the onset of symptoms in *Mecp2*-het mice is delayed and more variable than in males (25), with some females being asymptomatic even by 5 months of age (27), future studies should investigate pubertal development in a larger cohort of female mice, following their symptoms into older adulthood.

The hypothalamic GnRH neurons are key regulators of gonadal hormones and hence of pubertal onset and fertility. Previous reports have shown hypothalamic dysfunction in *Mecp2*-mutant mice, including in the hypothalamic-hypophyseal-adrenal axis (8,45); and in leptin pathways (46,47). Indeed, a recent report studying *Mecp2*-mutant mice showed that, whereas the growth of most brain areas (such as neocortex and hippocampus) is delayed in males and females, this growth is completely arrested in the hypothalamus, suggesting a more severe impairment of this important neuroendocrine structure (48). However, to our knowledge, our study is the first to report an increased density of GnRH neurons in the septo-hypothalamic region in *Mecp2^CD1^*-null males. Excess of hypothalamic GnRH neurons due to enhanced survival and migration was demonstrated following deletion of neuropilin-1 in GnRH neurons in a mouse model of central precocious puberty (49).

The increased number of GnRH neurons in *Mecp2^CD1^*-null males, however, did not lead to increased GnRH circulating content, but rather a slightly lower concentration of this hormone. In addition, we found significantly lower circulating levels of both LH and testosterone in *Mecp2^CD1^*-null males, in agreement with a previous study (28). Several mechanisms might explain this finding. As described, the low body weight of these males may have led to delayed puberty with hypogonadism. In circumstances of low nutrition and chronic disease, GnRH pulsatility is impaired, with consequent functional hypogonadotropic hypogonadism and thus delayed pubertal onset. It is conceivable that loss of *Mecp2* might result in increased hypothalamic GnRH neurons, but functional hypogonadism secondary to the whole-body effects of *Mecp2* deficiency might overcome this, meaning that instead of precocious puberty, a phenotype of delayed puberty is seen in this model. In keeping with this, it is interesting that *Mecp2^CD1^*-null males show pubertal onset at a lower body weight than healthy males.

In terms of the mechanism by which loss of *Mecp2* could affect GnRH neurons, this might occur through a dysregulation of Erα signaling, which is regulated by MeCP2 (24). The loss of *Mecp2* could also affect cells upstream of GnRH, for example, via arcuate nucleus kisspeptin modulation of GnRH release. As described, MECP2 colocalises with the histone mark H3K27me3. Enhanced H3K27me3 content at the 5′ regulatory region of the Kiss1 gene silences its expression (20), whilst reduction leads to increased *Kiss1* expression, GnRH pulsatility and pubertal development. Male mice with disrupted *Kiss1* expression show a reduction in testis size as well as lower levels of testosterone (50). Additionally, a study using a mouse model of *MECP2* duplication syndrome found that circulating levels of testosterone were increased in these mice (51). In these mice, it was shown that *Mecp2* was expressed in Leydig cells and led to an increase of androgen synthesis via an upregulation of the receptor for LH (LHCGR) and a reduction in aromatization to produce oestrogens (51). Although we did not find visible alterations in Leydig cells in our samples, future studies should address whether *Mecp2^CD1^*-null males display lower levels of LHR in these cells. Finally, the lower levels of LH found in these mice would contribute to a lesser stimulation of Leydig cells, leading to the observed lower levels of testosterone.

In female mice, however, we did not find significant differences in number of GnRH cells. As with pubertal timing, the later onset of symptoms in *Mecp2^CD1^*-het females might explain this difference. Further, since the MeCP2 protein is expressed in the nucleus of GnRH neurons, skewed X inactivation in these cells could contribute to different outcomes in terms of symptoms (52,53) and pubertal timing. Future studies assessing the GnRH system and circulating levels of GnRH, LH and sex steroids in older females might reveal relevant alterations.

The effects of pubertal dysregulation and altered hormonal signalling have an influence beyond the reproductive axis. Thus, brain circuits dependent on the levels of sex steroid hormones, such as the AVP-ergic system in males, display an aberrant configuration in *Mecp2*-null males, as seen previously (28) and in the present study. These circuits are involved in the control of emotional and social behaviour (54,55), highlighting the necessity of further investigating the mechanism by which *Mecp2* deficiency may impact their wiring, with the aim of finding novel therapeutic targets to potentially ameliorate emotional and social impairment in RTT and related conditions.

## Limitations

Despite the high validity of the model, there are still a number of concerns when trying to translate animal data to humans. For example, in humans, boys usually die perinatally, and females develop the first symptoms during early infancy, whereas in mice, males display an overt phenotype around adolescence and females after young adulthood (25,27). Hence, the biological age of the animals and humans in terms of symptomatology is not comparable. Second, the pubertal markers used in the mouse do not completely match those of humans. In spite of these limitations, our data in the mouse model supports a disruption of the hypothalamic-pituitary-gonadal axis caused by *Mecp2* deficiency, whose mechanism warrants further investigation.

## Conclusions

In this study, we found that a mouse model deficient for *Mecp2* display reduction in body weight during pubescence and dysregulated pubertal timing and function of the hypothalamic-pituitary-gonadal axis. The effects found are strongly significant in males, hemizygous for the mutation. By contrast, we did not find significant differences in females, which could be related to heterozygosity leading to variable symptom onset. Further, we found a significant increase in GnRH neurons but a decrease in circulating GnRH, LH and testosterone in males, suggesting a general malfunction of the hypothalamic-hypophyseal-gonadal axis. Since pubertal dysregulation has been observed in human patients with mutations in *MECP2*, our study can be used as a starting point to further investigate the biological mechanism by which *MECP2* dysfunction contributes to alterations in puberty.

## Statements

### Availability of data

The datasets used and/or analysed during the current study are available from the corresponding author on reasonable request.

## Ethics approval and consent to participate

All experimental procedures were approved by the Committee of Ethics and Animal Experimentation of the Universitat de València and treated according to the European Union Council Directive of June 3rd, 2010 (6106/1/10 REV1) and under an animal-usage license issued by the Direcció General de Producció Agrària i Ramaderia de la Generalitat Valenciana (2022/VSC PEA/0288).

## Permission to reproduce material from other sources

Not applicable

## Practical Applications

n/a

## Authors’ contributions

AMS performed experimental procedures, acquired images, analysed and interpreted data, prepared figures and wrote the manuscript; DJD, REP and AVT performed experiments and acquired images; JER contributed to data interpretation and writing; SRH obtained funding, designed the study, contributed to data interpretation and writing. CAP obtained funding, designed the study, performed and supervised experimental procedures, performed data analysis and wrote the manuscript. Final version of this manuscript was discussed and approved by all authors.

## List of abbreviations

AVP: Arginine-vasopressin
BPS: Balanopreputial separation
CPP: Central precocious puberty
DAPI: 4’,6– diamino-2-fe-niindol
ERβ: Estrogen receptor-β
FSH: Follicle stimulant hormone
FXYD1: FXYD domain-containing transport regulator 1
GnRH: Gonadotropin Realising Hormone
LH: Luteinizing hormone
LHb: Lateral habenula
LHR: Receptor for LH
MDS: MECP2 duplication syndrome
MECP2: Methyl CpG-binding protein 2
NDS: Normal donkey serum
Pa: Paraventricular nucleus
PBS: Phosphate saline solution
PFA: Parafolmaldehyde
RTT: Rett syndrome
TBS: TRIS buffered saline
VO: Vaginal opening

## Acknowledgements

Authors are indebted to Paloma Sevilla-Ferrer and Carmen Tejada Cortés for technical assistance, to the personnel from the Animal Production and Experimentation of the Central Service for Experimental Research (SCSIE, UV) for their help with mice husbandry, to Dr Jose V. Torres-Pérez and Dr María Abellán-Álvaro for their help to set up the puberty measures, and to Prof. Fernando Martínez García, Dr MªJosé Sánchez Catalán and Prof. Enrique Lanuza for discussion of results.

