## Supplementary material for "Pubertal development and hypothalamic-pituitary-gonadal axis are altered in male mice lacking *Mecp2*"

[§] Equal contribution

*Corresponding authors

***Protocol for Immunofluorescence for MeCP2 and GnRH protocol***

Double immunostaining for MeCP2 and GnRH was performed in one out of five parallel brain sets from WT and *Mecp2$^{CD1}$*-null males and MeCP2 immunodetection was used for one out of five parallel brain series.

Free-floating sections were washed three times with 0.05 M TBS pH 7.6 (TBS). In brief, sections were (i) previously treated with citrate buffer 0.01M pH 7.6 for 30 min at 80ºC. Then, sections were (ii) pre-incubated in 3% NDS in 0.05 M TRIS buffered saline pH 7.6 (TBS) with 0.3% Triton X-100, at RT for 1 h, to block nonspecific labelling; (iii) incubated in primary antibodies, rabbit anti-GnRH primary antibody (1:5000, Invitrogen, AB1567) and/or mouse anti-MeCP2 (Invitrogen, MA5-33096; 1:1000, previously used in (1,2)) diluted in TBS with 0.3% Triton X-100 with 4% NDS at 4ºC for 48 h; (iv) incubated with fluorescent-labelled secondary antibodies (Alexa Fluor 488 donkey anti-mouse 1:400; Invitrogen A21202 and/or Rhodamine Red-X donkey anti-rabbit 1:400; Jackson ImmunoResearch, 711- 295-152) diluted in TBS for 2 h at RT. (v) To reveal the cytoarchitecture in brain sections, they were counterstained prior to mounting by bathing them for 1 min in DAPI (a nuclear staining) at RT. After each step, sections were washed three times for 5 min in TBS except between steps (ii) and (iii). Finally, sections were washed in TB, mounted onto gelatinised slides and cover-slipped with fluorescence mounting medium FluorSave Reagent (Sigma-Aldrich, 345789).

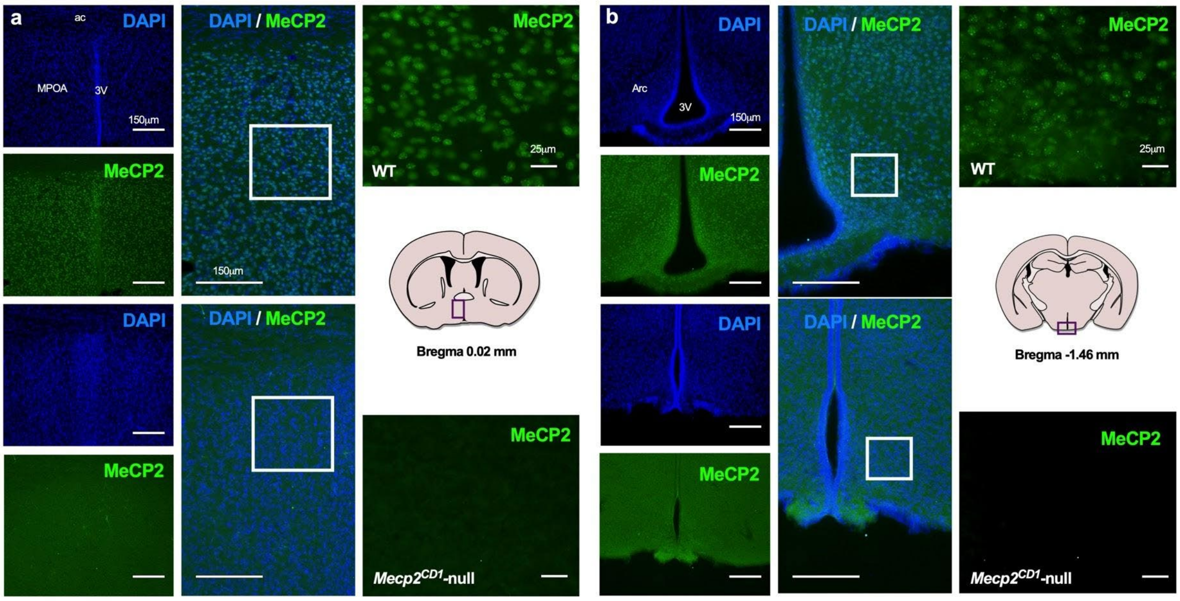

**Figure S1. MeCP2 is expressed in the hypothalamus of WT mice, but not in *Mecp2*$^{CD1}$-null brain tissue.** Representative images of MeCP2-containing nuclei (green) and DAPI (blue) in the medial preoptic area (MPOA, a; Bregma 0.02mm) and arcuate nucleus (Arc, b; Bregma -1.46mm) in WT and *Mecp2*$^{CD1}$-null mice. MeCP2 expression is completely absent in *Mecp2*$^{CD1}$-null brain samples in comparison to WT, in which most of the DAPI-positive nuclei co-localize with MeCP2.

***Breeding capacity of Mecp2-het females***

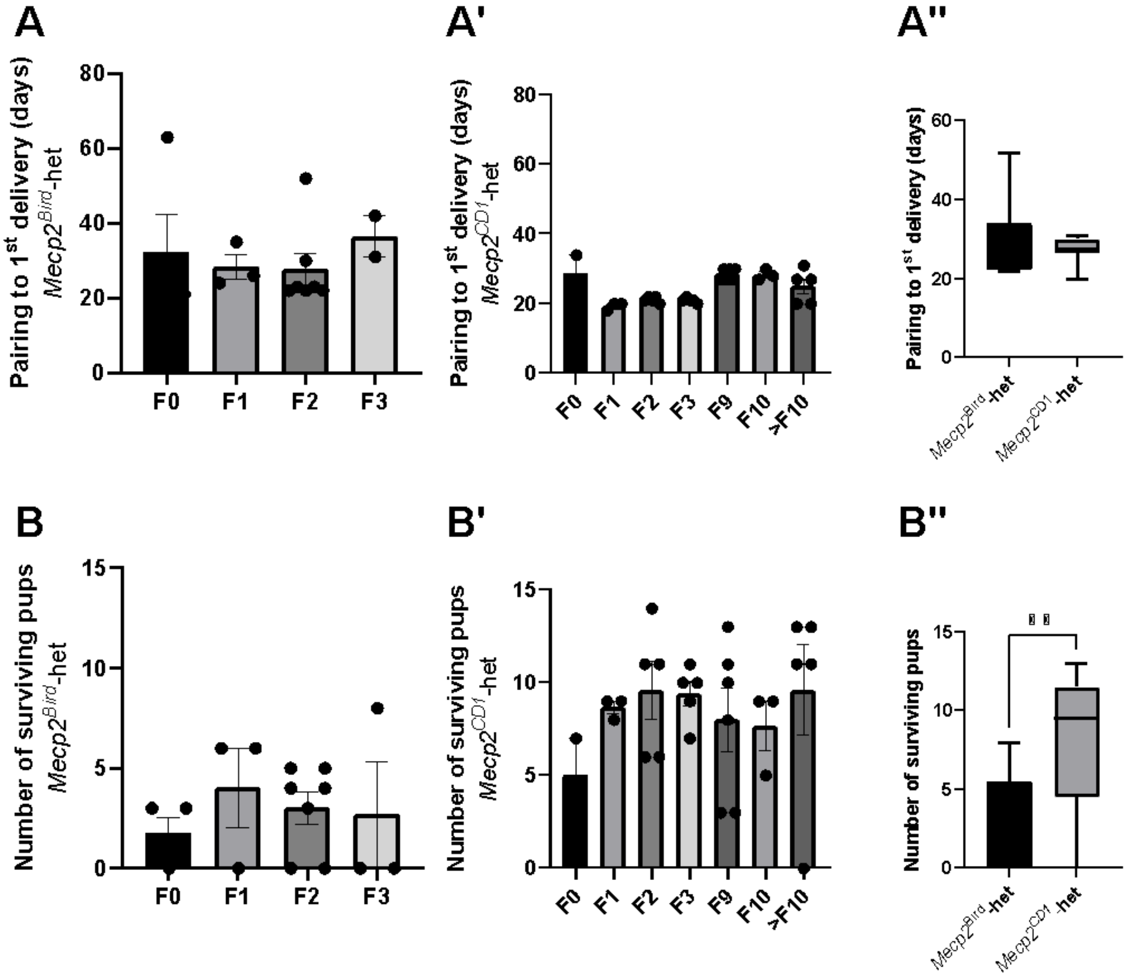


**Figure S2. Derivation to CD1 background increases colony productivity.** Graphs showing the number of days elapsed from pairing with a stud male until first delivery in *Mecp2*$^{Bird}$-het females (A) and *Mecp2*$^{CD1}$-het females (A'), across generations. In the case of *Mecp2*$^{Bird}$-het, F0 represent the first crossing of 4 females purchased to the Jackson Lab; F1, F2 and F3 are the in-house successive crossings. In the case of *Mecp2*$^{CD1}$-het females, F0 represents the first crossing of 2 *Mecp2*$^{Bird}$-het bred in house with 2 CD1 stud males. The successive F1…Fn crossings have been performed in house in a 2 females x 1 male scheme. To analyse the effect of strain in this measure, we compared the time elapsed from pairing to first delivery in F1-F3 in *Mecp2*$^{Bird}$-het (n=13) to F9->F10 in *Mecp2*$^{CD1}$-het (n=14) (A''), and found no significant effect of strain. By contrast, strain had a significant effect in the number of pups surviving until weaning. Graphs show this measure in *Mecp2*$^{Bird}$-het (A), *Mecp2*$^{CD1}$-het (A'), and a the comparison between F1-F3 of *Mecp2*$^{Bird}$-het and F9->F10 in *Mecp2*$^{CD1}$-het (A''). **, $p < 0.01$, Mann-Whitney test.

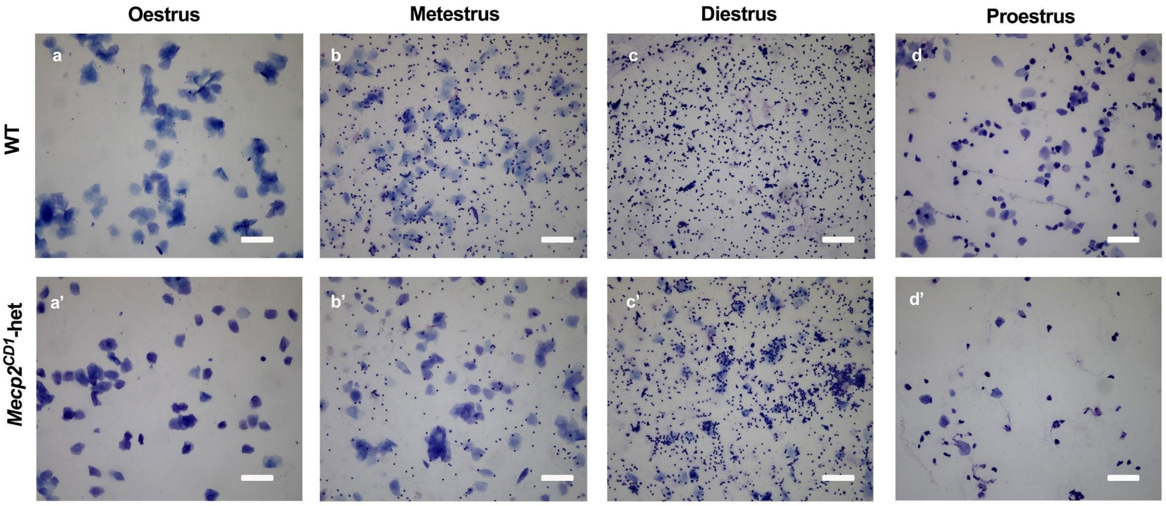


**Figure S3. Photomicrographs of toluidine blue-stained vaginal smears from *Mecp2$^{CD1}$*-het and WT females at different phases of the oestrus cycle.** a, a') Oestrus, showing almost exclusively cornified cells; (b, b') metestrus is characterised by the presence of a mix of cell types, mostly cornified cells and leukocytes, which are predominant in (c,c') diestrus phase and (d,d') proestrus, showing a high proportion of nucleated epithelial cells. Scale bar: 50μm.

*Immunofluorescence for GnRH and MeCP2*

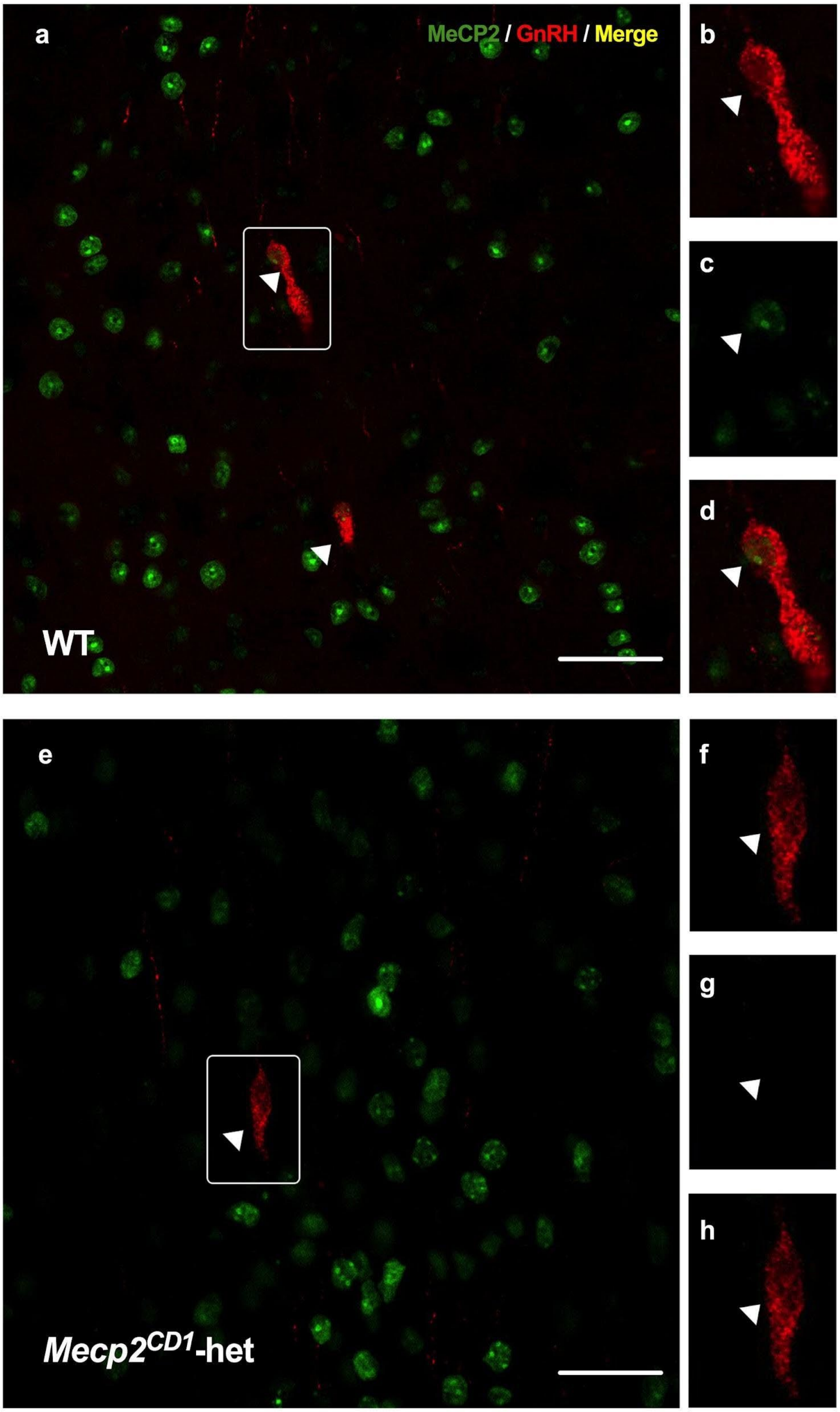

**Figure S4. Co-localization of MeCP2 and GnRH-ir neurons in septo-hypothalamic area.** Representative single confocal plane of MeCP2-containing (green; c and g) and GnRH-containing (red; b and f) neurons. Noteworthy that MeCP2 signalling co-localizes with GnRH (a and d) in WT female septo-hypothalamic area (Bregma 0.5-0.02 mm), whereas the GnRH-ir cell did not express MeCP2 (e and h) in *Mecp2*$^{CD1}$-het female. Scale bar: 50μm.

***Arginine vasopressin immunohistochemistry***

We performed a pilot study by immunostaining for AVP in a subset of one of five parallel sets (WT, n = 3; *Mecp2$^{CD1}$*-null, n = 3) as previously published (3). Briefly, sections were washed three times with 0.05 M TBS for 5 min. Then, they were incubated sequentially in: (i) 1% $H_2O_2$ in 0.05 M TBS pH 7.6 for 30 min at RT for endogenous peroxidase inactivation; (ii) blocking solution, 0.05 M TBS pH 7.6 with 0.3% Triton X-100 and 2% normal goat serum; (iii) primary antibody (1:10,000, rabbit anti-vasopressin IgG, Chemicon, AB1565) overnight at 4 °C; (iv) diluted biotinylated secondary antibody (1:200, goat anti-rabbit IgG, Vector Labs, BA-1000) in TBS for 90 min at RT; (v) avidin–biotin–peroxidase complex (ABC Elite kit; Vector Labs, PK-6200) in TBS for 90 min at RT. Between each step, sections were washed in TBS (3 × 10 min) except after step (ii). After ABC incubation, sections were rinsed in TBS (3 × 10 min) and TRIS buffer (TB) 0.05 M, pH 8 (3 × 10 min). The histochemical detection of the resulting peroxidase activity was performed by incubation in 0.003% $H_2O_2$ and 0.025% 3,3-diaminobenzidine (Sigma) in TB for about 15 min. The sections were finally rinsed thoroughly in TB, mounted onto gelatinized slides, dehydrated in ethanol, cleared with xylene and coverslipped with Entellan.

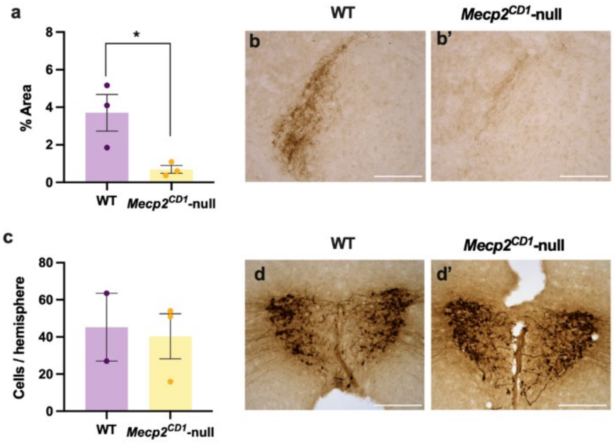


**Figure S5. Lack of *Mecp2* reduces testosterone-dependent AVP-ergic innervation in habenula in *Mecp2$^{CD1}$*-null males without affecting AVP-ergic neurons in the paraventricular nucleus of the hypothalamus.** Testosterone-dependent AVP-ergic innervation in habenula in *Mecp2$^{CD1}$*-null males is significantly reduced in comparison to WT animals (a, b and b'). By contrast, the density of AVPergic cells in the paraventricular nucleus of the hypothalamus is not affected by genotype (c, d, d'). Data are shown as Mean ± S.E.M. Student's t-test, * $p < 0.05$. Scale bar: 50μm
